# Towards the quantitative characterization of piglets’ robustness to weaning: A modelling approach

**DOI:** 10.1101/428920

**Authors:** M. Revilla, N.C. Friggens, L.-P. Broudiscou, G. Lemonnier, F. Blanc, L. Ravon, M.-J. Mercat, Y. Billon, C. Rogel-Gaillard, N. Le Floch, J. Estellé, R. Muñoz-Tamayo

## Abstract

Weaning is a critical transition phase in swine production in which piglets must cope with different stressors that may affect their health. During this period, the prophylactic use of antibiotics is still frequent to limit piglet morbidity, which raises both economic and public health concerns such as the appearance of antimicrobial-resistant microbes. With the interest of developing tools for assisting health and management decisions around weaning, it is key to provide robustness indexes that inform on the animals capacity to endure the challenges associated to weaning. This work aimed at developing a modelling approach for facilitating the quantification of piglet resilience to weaning. We monitored 325 Large White pigs weaned at 28 days of age and further housed and fed conventionally during the post-weaning period without antibiotic administration. Body weight and diarrhoea scores were recorded before and after weaning, and blood was sampled at weaning and one week later for collecting haematological data. We constructed a dynamic model based on the Gompertz-Makeham law to describe live weight trajectories during the first 75 days after weaning following the rationale that the animal response is partitioned in two time windows (a perturbation and a recovery window). Model calibration was performed for each animal. Our results show that the transition time between the two time windows, as well as the weight trajectories are characteristic for each individual. The model captured the weight dynamics of animals at different degrees of perturbation, with an average coefficient of determination of 0.99, and a concordance correlation coefficient of 0.99. The utility of the model is that it provides biological parameters that inform on the amplitude and length of perturbation, and the rate of animal recovery. Our rationale is that the dynamics of weight inform on the capability of the animal to cope with the weaning disturbance. Indeed, there were significant correlations between model parameters and individual diarrhoea scores and haematological traits. Overall, the parameters of our model can be useful for constructing weaning robustness indexes by using exclusively the growth curves. We foresee that this modelling approach will provide a step forward in the quantitative characterization of robustness.

**Implications:** The quantitative characterization of animal robustness at weaning is a key step for management strategies to improve health and welfare. This characterization is also instrumental for the further design of selection strategies for productivity and robustness. Within a precision livestock farming optic, this study develops a mathematical modelling approach to describe the body weight of piglets from weaning with the rationale that weight trajectories provide central information to quantify the capability of the animal to cope with the weaning disturbance.

## Introduction

In modern swine breeding conditions weaning is one of the most critical phases (Lallès *et al*., 2007). In industrial farming this usually occurs at around 3-4 weeks of age, although natural weaning occurs around 17 weeks after birth (Jensen *et al*., 1986). Weaning can be defined as a sudden, short, and complex event characterized by changes in diet, social, and environmental conditions (Campbell *et al*., 2013). The switch from highly digestible liquid milk to a less-digestible and more complex solid feed has consequences on the physiology of the gastrointestinal tract, causing a transitory anorexia, intestinal inflammation, and unbalanced gut microbiota (Pié *et al*., 2014). Indeed, these changes trigger the development of a dysbiotic state of the gut microbiota that can ultimately result in enteric disease and diarrhoea (Gresse *et al*., 2017). In general, it has been estimated that in commercial conditions the first day after weaning piglets lose about 100-250 g body weight (BW) regardless of weaning age (LeDividich and Sève, 2000). For this reason, during this period, the prophylactic use of antibiotics is still frequent to limit piglet morbidity and diarrhoea episodes. However, this procedure raises economic and public health concerns such as the growing number of antimicrobial-resistant agents. In this context, finding antibiotic alternatives to maintain piglet health at the critical weaning period and preserve public health becomes a real emergency. Consequently, there is an increased interest to develop tools for assisting health (European Medicines Agency, 2017) and management decisions around this critical period.

The response of piglets to weaning relates to its robustness, that is its capacity to maintain productivity in a wide range of environments without compromising reproduction, health, and welfare (Friggens *et al*., 2017). A key component of robustness, sometimes termed resilience, is the ability to cope with environmental perturbations. One way to characterize resilience is by quantifying the extent of deviations from the non-perturbed trajectories of physiological functions (Codrea *et al*., 2011). In this respect, the development of mathematical models in animal science can be of help to gaining insight in animal robustness (e.g., Sadoul *et al*., 2015). The Gompertz model (Gompertz, 1825) is well known and widely used to describe the growth of animals. However, the Gompertz model does not account for the effects caused by a perturbation. To consider the effect of a disturbing environment, William Makeham extended the Gompertz model by adding a constant term to explain that the rate of change is also driven by factors that are independent of the age (Makeham, 1873). The resulting equation is known as the generalised Gompertz-Makeham equation (Golubev, 2009). Based on this scenario, the aim of the present study was to develop a modelling approach for facilitating the quantification of piglet resilience at weaning. We developed a perturbed model to describe animal growth based on the Gompertz-Makeham equation. The model parameters have biological interpretation and inform on the amplitude and length of the perturbation. After defining these biological parameters, we performed correlation analyses with faeces score data and haematological traits in order to evaluate the pertinence of these parameters for assessing differences in robustness.

## Material and methods

### Animal samples

The animal resource population used in this study comes from *Institut National de la Recherche Agronomique* (INRA)’s experimental farm (*Le Magneraud*, France). Here, we report results based on 325 piglets from a Large White selected line within the Pigletbiota ANR project (http://www.agence-nationale-recherche.fr/Project-ANR-14-CE18-0004). All animal procedures were performed according to the guidelines for the care and use of experimental animals established by INRA (Ability for animal experimentation to J. Estellé: R-94ENVA-F1-12; agreement for experimentation of INRA’s *Le Magneraud* farm: A17661; protocol approved by the French Ministry of Research with authorization ID APAFIS#2073-2015092310063992 v2 after the review of ethics committee n°084).

Piglets were weaned at an average age of 28.7 ± 1.6 days (± standard error of the means; SEM) and weighted 8.91 ± 0.49 kg. All animals were maintained under standard intensive conditions and feeding was *ad libitum* with a cereal-based standard diet. The management, environmental and housing factors were controlled for all animals during the whole study. None of the animals received antibiotic treatments during the study, and were free of the principal swine infectious agents.

### Measures of animal’s body weight

BW data were collected at birth, then three times before the weaning period (once per week), two times per week from 29 to 50 days, once per week from 50 to 100 days, and then once every two weeks until the end of the animal’s life. On average, each pig was weighted 20 times.

### Faeces score data

Faeces were individually observed at days 0, 2, 6, 8, 12, 15, 20, 27, and 34 with respect to the weaning, and were scored in three levels being: 0 for normal, 1 for soft faeces but without diarrhoea, and 2 for evident diarrhoea (Le Floc’h *et al*., 2014). The faeces score data was analyzed in three different ways: (1) by summing the number of diarrhoea measurements and correcting by the number of observations per animal (FS_sum); (2) by group levels, being level 0 for those animals that have no diarrhoea observations on any of the measures, level 1 for those that have only one diarrhoea record, and level 2 for the animals that have shown two or more diarrhoea records (FS_gr); and (3) by presence or absence of diarrhoea records (FS_p_a) taking into account all the observations made.

### Blood sampling and haematological traits

Blood samples were collected at 28 and 34 days of age from the jugular vein just before feeding time in Vacutainer tubes of 4 mL containing EDTA as anticoagulant. The haematological analyses were carried out in the following hours after sampling, using a MS9-5 instrument (Melet Schloesing; France). The hemograms included the following measures: Basophiles (Bas) [%], Blood plate (Plt) [m/mm3], Eosinophils (Eos) [%], Erythrocytes (Ery) [m/mm3], Hematocrit (Hct) [%], Hemoglobin (Hgb) [g/dl], Leucocytes (Leu) [m/mm3], Lymphocytes (Lym) [%], Mean corpuscular hemoglobin concentration (MCHC) [g/dl], Mean corpuscular haemoglobin content (MCH) [pg], Mean corpuscular volume (MCV), Monocytes (Mon) [%], and Neutrophils (N) [%]. In addition, the Neutrophils/Lymphocyte ratio (Ratio N/Lym) [%] was estimated as a measure of stress (Puppe, 1997).

### Mathematical modelling of growth with the Gompertz function

The global dynamics of many natural processes including growth have been described by the Gompertz function (Waliszewski and Konarski, 2005). The essential characteristic of this function is that it exhibits an exponential decay of relative growth rate, making this model a reference for describing many types of growth of organisms. As first step for quantifying robustness, we focused on the dynamics of non perturbed live weight during the first 75 days after weaning using the Gompertz equation (1) using the formulation of Schulin-Zeuthen *et al*. (2008).

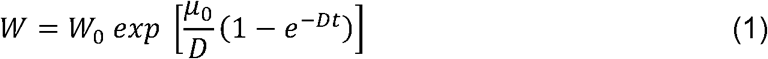

Where *W_0_* is the initial value of live weight *W* (kg), *μ_0_* is the initial value of the specific growth rate *μ* (d^−1^), the constant *D* (d^−1^) is a growth rate coefficient that controls the slope of the growth rate curve, and *t* (d) is time since weaning.

### Perturbed growth modelling with the Gompertz-Makeham function

The Gompertz model is a monotonic function that does not account for possible decrease of weight gain due to perturbations. To elaborate further on our hypothesis to quantify robustness at weaning period, we took as basis the Gompertz-Makeham extension (Golubev, 2009) that has the advantage of including explicitly the situation of the perturbation and thus allows to describe weight decrease. The weight dynamics is described by the following model with two ordinary differential equations

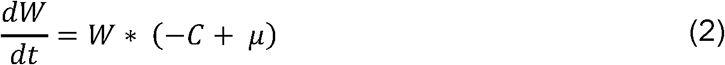

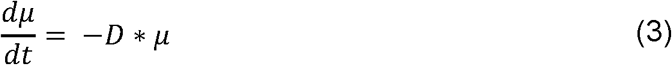

where *C* (d^−1^) is a parameter representing the effect of the environment on the weight change. Note that if *C* = 0, the equations (2-3) are the differential form of the classic Gompertz equation. Equations (2-3) can be arranged into one equation.

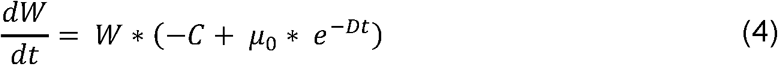

In the remaining of the text, we refer to the model in Eq. (4) as perturbed model. The specificity of our approach relies on the hypothesis that the weight dynamics of the animal is partitioned in two time windows (perturbed and recovery windows) to represent the moment at which the animal is perturbed and the moment at which it recovers from the perturbation. This is translated mathematically by modulating the parameter *C* with the following conditions

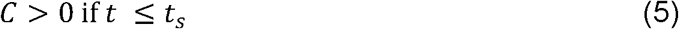

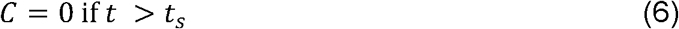

where *t_s_* is the time of the recovery switch that is assumed to be specific for each animal.

### Model evaluation and calibration

We tested the structural identifiability of both the Gompertz and the perturbed models. That is, we determined if it was theoretically possible to determine uniquely the model parameters given the available measurements (see; e.g., (Muñoz-Tamayo *et al*., 2018) for a discussion on structural identifiability). Identifiability testing was performed using the freely available software DAISY (Bellu *et al*., 2007). Both models are structurally globally identifiable, implying that the parameter estimation problem is well-posed (it has unique solution).

The mathematical models (Gompertz and perturbed) were further calibrated for each animal by minimizing the least squares error:

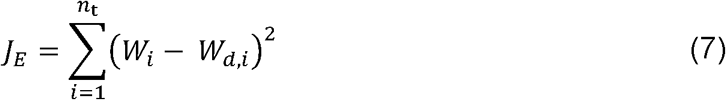

where *W_d_* is the weight data (kg), *W* the weight predicted by the model, and *n_t_* the total number of measurements. The numerical estimation of the parameters was performed in Scilab using the Nelder-Mead algorithm implemented in the *“fminsearch”* function (Scilab Enterprises, 2012) (v.6.0.0).

To allow comparison between all animals within the studied population, for each individual the objective function was further weighted with respect to the number of measurements for each animal (*n_t_*).

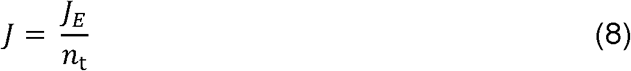

In addition, we carried out a calibration of the Gompertz model using the BW at weaning and only the last four records during the first 75 days after weaning of each animal (*n*_t_ = 5). We assumed that the resulting model calibrated with these data is an approximation of the theoretical growth rate of the animals not experiencing any perturbation. In the remaining text, we will call the trajectory resulted from this calibration with the subset data as the unperturbed curve.

The Akaike Information Criterion (AIC) (Akaike, 1974) was used to select the best candidate model. The AIC is a trade-off between goodness of fitness and model complexity. The AIC was calculated as follows (Banks and Joyner, 2017).

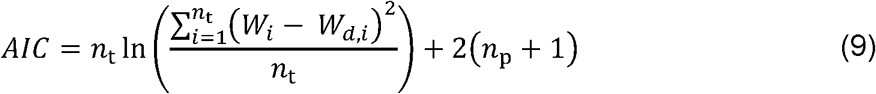

where *n*_p_ is the total number of model parameters. When there are several competing models, the best model in the AIC sense is the one with the smallest AIC.

Model performance was assessed by classical statistical indicators, namely coefficient of determination (r^2^) and the Lin’s concordance correlation coefficient (CCC) (Lin, 1989).

### Statistical analyses to relate model parameters with haematological traits

Correlation analyses were performed to explore the relationships between the perturbed model parameters and the faeces score data and haematological indicators measured in the blood formula. Pearson and Spearman correlations among them were analyzed in R using the “*cor*” function in the base package (R Core Team, 2017). When required, data was normalized applying the log_2_ or log_10_ transformation. Visualization of correlations was performed with the *“corrplot”* R package (Wei and Simko, 2017). Only the correlations with *p*-value less than 0.05 were considered as significant and were thus represented.

## Results

### Growth curve modelling from weaning

To get an overview of the piglets’ response to weaning, the BW measurements of the first 75 days after this critical period were analyzed. Figure 1 displays the BW dynamic trajectories of four animals exhibiting two extreme patterns. The experimental data is compared to predicted responses given by both the Gompertz and perturbed model. An animal ranking was performed with respect to the goodness of fitting for the Gompertz equation given by the criterion *J* in Eq. (8). This ranking provided a first indication on the magnitude of the perturbation and animal resilience, *i.e* the higher *J* the higher the level of perturbation (Supplementary Table S1). Moreover, the analysis of *J* reflects a difference of one order of magnitude between the strongest perturbed animal and the less perturbed one. As observed in Figure 1, the perturbed model captured accurately the weight dynamics of animals at different degrees of perturbation (Table 1; Supplementary Table S1). The minimum and the mean values of the r^2^ coefficient were 0.62 and 0.99, respectively. As well, for the CCC coefficient the minimum value was 0.73, and the mean 0.99 (Supplementary Table S1).

**Figure 1.**
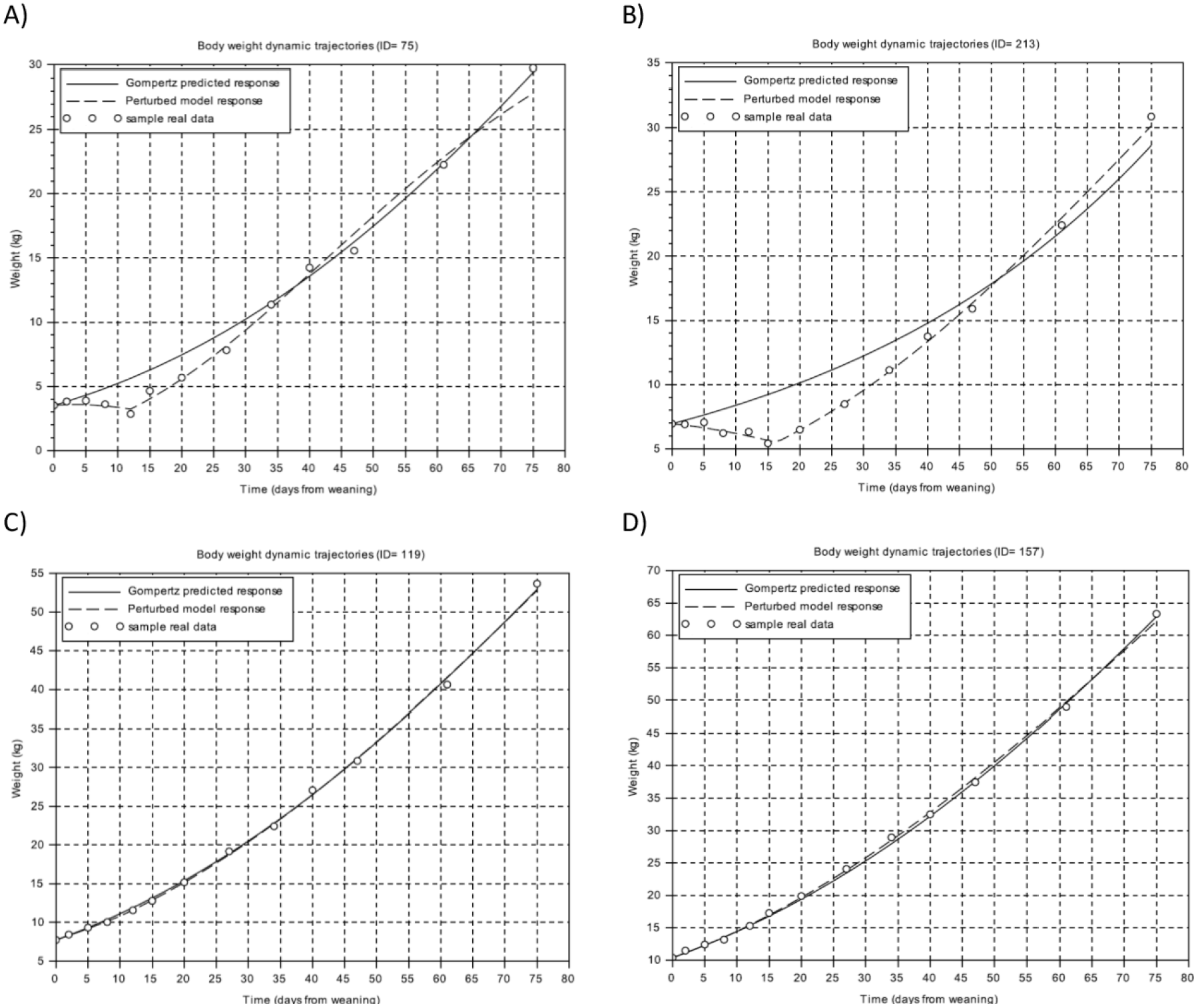
Body weight dynamics trajectories. A and B figures represent samples with the worst level of fitting using the Gompertz model. C and D figures represent samples with the best fitting using the Gompertz model. Circles represent the different body weight measures of the individual piglet relative to days from weaning, the solid line is the Gompertz predicted response, and the dashed line is the perturbed model response.

**Table 1.** Descriptive statistics for the parameters of the perturbed model.

| Perturbed model parameters | Range | Mean | Std. Deviation |
| --- | --- | --- | --- |
| $\mu_0$ | $1.22 \times 10^{-02} - 1.11 \times 10^{-01}$ | $4.78 \times 10^{-02}$ | $1.22 \times 10^{-02}$ |
| $D$ | $1.24 \times 10^{-03} - 4.25 \times 10^{-02}$ | $1.53 \times 10^{-02}$ | $5.57 \times 10^{-03}$ |
| $C$ | $4.97 \times 10^{-10} - 1.00 \times 10^{-01}$ | $2.84 \times 10^{-02}$ | $1.47 \times 10^{-02}$ |
| $t_s$ | 0.00 - 23.75 | 9.94 | 3.41 |
| $J$ | $3.70 \times 10^{-05} - 5.42 \times 10^{-03}$ | $1.00 \times 10^{-03}$ | $7.79 \times 10^{-04}$ |
| $ABC$ | 0.00 - 137.86 | 29.72 | 22.96 |

$\mu_0$ = individual growth rate at the moment of weaning; $D$ = extent of the exponential decay of the growth; $C$ = level of perturbation; $t_s$ = moment at which the animal recover for the perturbation; $J$ = model error; $ABC$ = area between curves

According to the AIC criterion, the perturbed model is superior to the Gompertz model. In addition, the advantage of the perturbed model is that it provides biological parameters that inform on the amplitude and length of perturbation, and the rate of animal recovery. The parameter *μ_0_* characterizes the individual growth rate at the moment of weaning. The parameter *C* determines the level of perturbation: the higher the value of this parameter, the higher the degree of perturbation of the animal. The parameter *t_s_* indicates the moment at which the animal starts to recover from the perturbation. Finally, the parameter *D* indicates the extent of the exponential decay of the growth. The descriptive statistics and the estimated parameter values of the perturbed model are given in Table 1 and Supplementary Table S1, respectively. From the unperturbed curve for each animal obtained from the calibration of the Gompertz model using the BW at weaning and the last four records during the first 75 days after weaning of each animal, we calculated the area between the unperturbed curve and the perturbed curve from the time where weaning starts to the time where the curves intersect. The resulting value was called the Area Between Curves (*ABC*) index. It reflects the difference between the unperturbed curve and the perturbed model response (Figure 2) (Supplementary Table S1).

**Figure 2.**
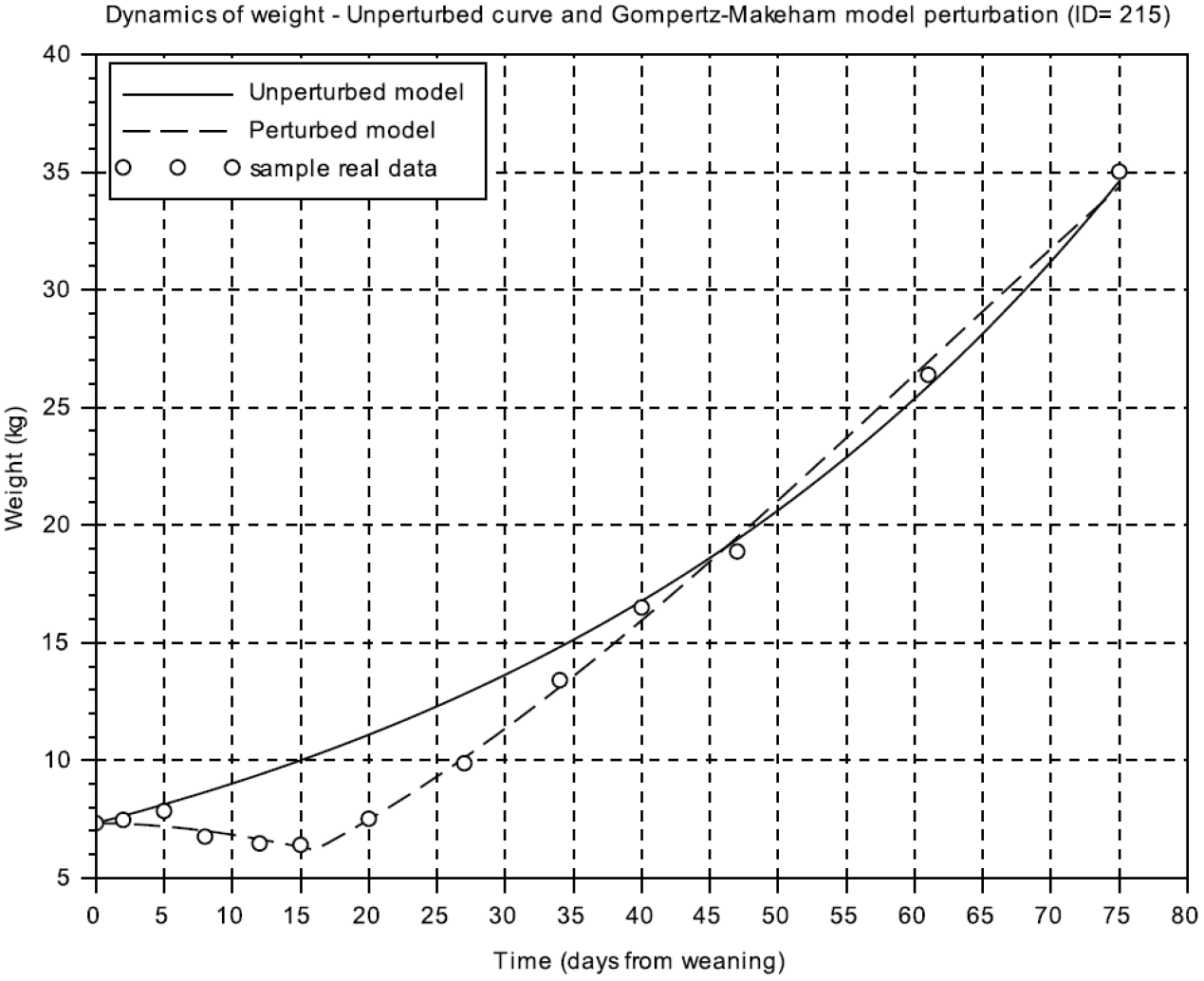
Comparison of the weight dynamics as predicted by the unperturbed and the Gompertz-Makeman (perturbed) models. Animal ID= 215 is represented. Circles represent the different body weight measures of the individual piglet relative to days from weaning, the solid line is the predicted response of the unperturbed model, and the dashed line is the perturbed model response.

The parameter *μ_0_* showed significant positive correlations with parameter *D* (r= 0.91, *p*-value= 5.78 × 10^−23^), parameter *C* (r= 0.60, *p*-value= 4.07 × 10^−23^), parameter *t_s_* (r= 0.39, *p*-value= 4.91 × 10^−13^), and parameter *ABC* (r= 0.29, *p*-value= 1.27 × 10^−07^). A moderate-to-strong significant positive correlation was estimated between the parameters *C* and *ABC* of the perturbed model (r= 0.64, *p*-value= 3.34 × 10^−17^) (Figure 3). There is an obvious correlation for those animals that have a greater degree of perturbation to be associated with a higher *ABC* parameter. In fact, the parameter *C* is a measure of the deviation between the unperturbed and the perturbed curves. A moderate positive correlation value was found between the parameter *C* and parameter *D* (r= 0.50, *p*-value= 5.07 × 10^−22^). In contrast, no significant correlation between parameter *C* and parameter *t_s_* was identified. Moreover, parameter *D* showed a positive correlation with the parameter *t_s_* (r= 0.31, *p*-value= 1.39 × 10^−8^). The weakest significant positive correlation was observed between parameter *D* and parameter *ABC* (r= 0.24, *p*-value= 1.66 × 10^−5^), and parameter *t_s_* and parameter *ABC* (r= 0.18, *p*-value= 1.44 × 10^−3^).

**Figure 3.**
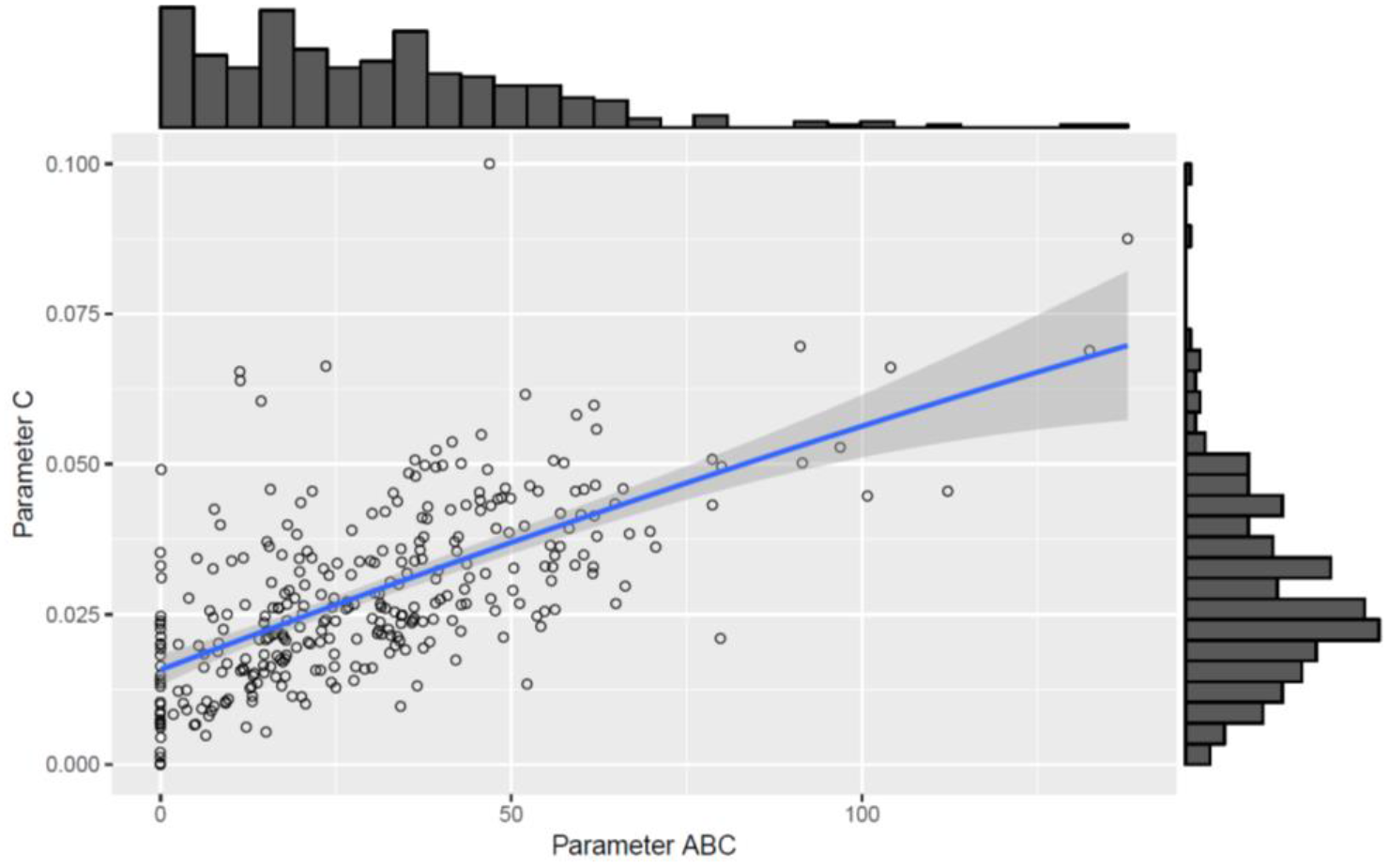
Scatter plot with marginal histograms illustrating the relationship between parameter *C* and parameter *ABC* of the Gompertz-Makeham perturbed model.

### Post-weaning diarrhoea scores and model parameters

The relationship between the model parameters and the faeces score data of weaned piglets were analyzed by correlation analyses (Figure 4). Several significant positive correlations were identified. Faecal score data, analyzed as a continuous variable (FS_Sum), was positively correlated to the parameter *C* (r= 0.29, *p*-value= 7.55 × 10^−8^), parameter *μ_0_* (0.22, *p*-value= 9.23 × 10^−5^), parameter *D* (r= 0.17, *p*-value= 1.90 × 10^−3^), and parameter *ABC* (r= 0.16, *p*-value= 4.35 × 10^−3^). Moreover, the faeces score data analyzed by groups (FS_gr; level 0= no diarrhoea observations; level 1= one diarrhoea record; level 2= two or more diarrhoea records) and the parameter *C*, parameter *μ_0_*, and parameter *D* have a correlation coefficient of 0.25 (*p*-value= 4.68 × 10^−6^), 0.17 (*p*-value= 2.64 × 10^−3^), and 0.12 (*p*-value= 2.66 × 10^−2^), respectively. The presence/absence of diarrhoea records shows significant correlations with parameter C (r= 0.18, *p*-value= 1.19 × 10^−3^), and parameter *μ_0_* (r= 0.12, *p*-value= 3.06 × 10^−2^).

**Figure 4.**
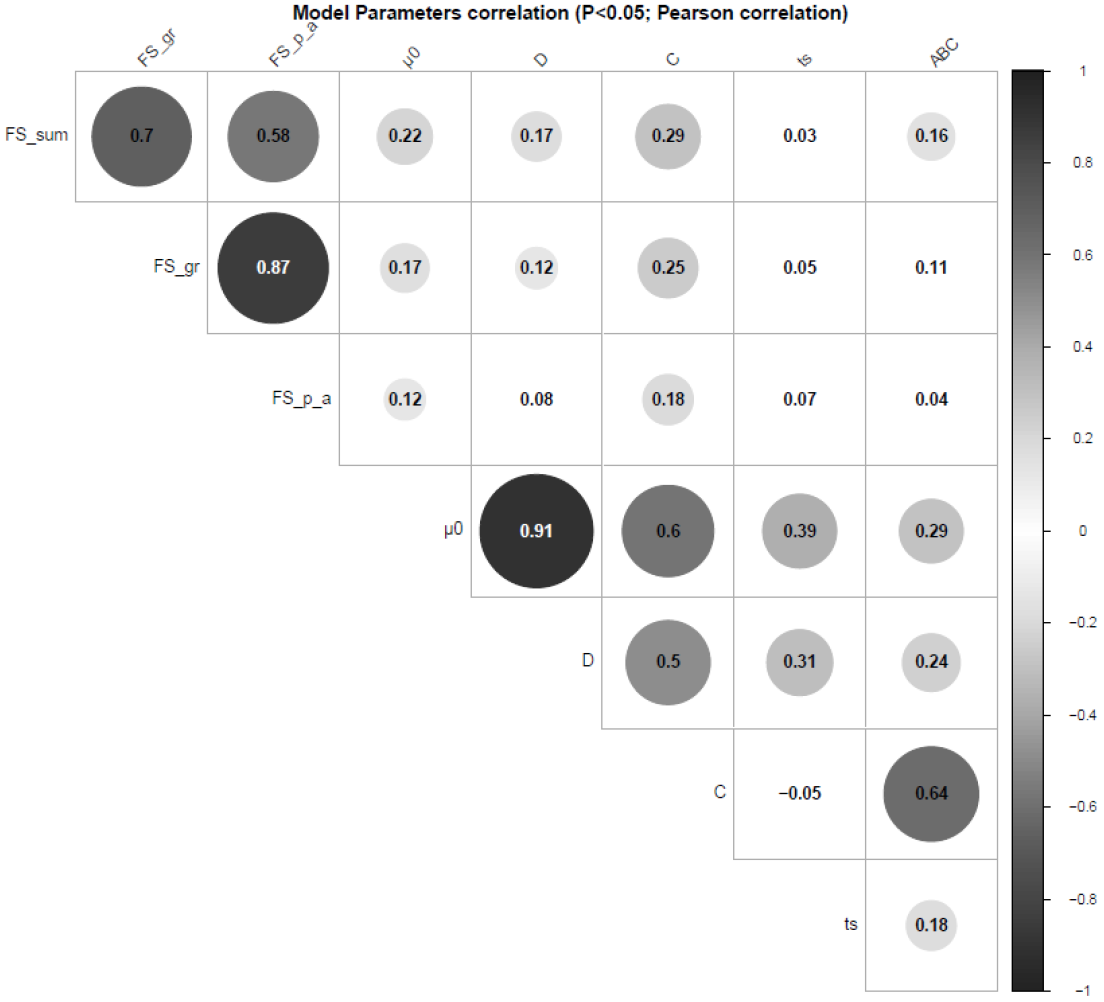
Pearson’s coefficients to visualize correlations among the model parameters of the Gompertz-Makeham perturbed model and the faeces score data. The size of the circles are proportional to the correlation coefficients. Only the correlations with *p*-value less than 0.05 were considered as significant and were represented with circles, the insignificant correlations are left blank.

### Blood cell population and model parameters

To evaluate piglet’s health status at weaning period, several blood cells measurements were recorded at weaning period (28 d) and one week later (34 d). The correlation analyses performed between the available haematological measurements of the 320 animals at 34 d and the model parameters showed positive and negative significant correlations (Table 2).

The results of the correlation analyses revealed the highest significant correlations between the parameter *D* and the Hgb (r= −0.23, *p*-value= 3.99 × 10^−5^), MCH (r= − 0.22, *p*-value= 6.75 × 10^−5^), and Plt (r= 0.22, *p*-value= 6.75 × 10^−5^).

The conducted correlation analyses display also moderate significant correlations between the developed parameter *ABC* and the rate of Mon (r= −0.30, *p*-value= 5.15 × 10^−8^) and Eos (r= 0.20, *p*-value= 2.54 × 10^−4^).

**Table 2.** Pearson’s coefficients to visualize correlations among the model parameters of the Gompertz-Makeham perturbed model and the haematological measurements (34 d) (N= 320 animals)

|  | *D* | *C* | *t<sub>s</sub>* | *ABC* | BW_w | Age_w | Leu | Lym | Mon | N | Eos | Bas | Ery | MCV | Hct | MCH | MCHC | Hgb | Plt | N/Lym |
| --- | --- | --- | --- | --- | --- | --- | --- | --- | --- | --- | --- | --- | --- | --- | --- | --- | --- | --- | --- | --- |
| $\mu_0$ | 0.90<sup>***</sup> | 0.52<sup>***</sup> | 0.43<sup>***</sup> | 0.30<sup>***</sup> | -0.29<sup>***</sup> | 0.19<sup>***</sup> | 0.03 | -0.12<sup>*</sup> | -0.02 | 0.13<sup>*</sup> | 0.11 | -0.04 | -0.03 | -0.05 | -0.06 | -0.11 | -0.15<sup>**</sup> | -0.12<sup>*</sup> | 0.16<sup>**</sup> | 0.11 |
| *D* |  | 0.43<sup>***</sup> | 0.40<sup>***</sup> | 0.24<sup>***</sup> | -0.05 | 0.25<sup>***</sup> | 0.07 | -0.09 | -0.08 | 0.10 | 0.15<sup>**</sup> | 0.01 | -0.03 | -0.18<sup>***</sup> | -0.18<sup>**</sup> | -0.22<sup>***</sup> | -0.15<sup>**</sup> | -0.23<sup>***</sup> | 0.22<sup>***</sup> | 0.09 |
| *C* |  |  | 0.00 | 0.57<sup>***</sup> | 0.03 | 0.13<sup>*</sup> | -0.12<sup>*</sup> | 0.01 | -0.12<sup>*</sup> | 0.02 | 0.13<sup>*</sup> | -0.10 | 0.14<sup>*</sup> | -0.16<sup>**</sup> | -0.03 | -0.18<sup>**</sup> | -0.09 | -0.07 | 0.13<sup>*</sup> | -0.01 |
| *t<sub>s</sub>* |  |  |  | 0.25<sup>***</sup> | -0.15<sup>**</sup> | 0.13<sup>*</sup> | -0.16<sup>**</sup> | 0.00 | -0.11 | 0.02 | 0.08 | -0.08 | 0.04 | -0.05 | -0.02 | -0.07 | -0.05 | -0.04 | 0.06 | 0.03 |
| *ABC* |  |  |  |  | 0.22<sup>***</sup> | 0.24<sup>***</sup> | -0.11<sup>*</sup> | -0.02 | -0.30<sup>***</sup> | 0.08 | 0.20<sup>***</sup> | -0.16<sup>**</sup> | 0.15<sup>**</sup> | -0.18<sup>***</sup> | -0.04 | -0.16<sup>**</sup> | -0.01 | -0.05 | 0.03 | 0.04 |

$\mu_0$ = individual growth rate at the moment of weaning; *D* = extent of the exponential decay of the growth; *C* = level of perturbation; *t<sub>s</sub>* = moment at which the animal recover for the perturbation; *ABC* = area between curves; BW\_w = body weight at weaning; Age\_w = age at weaning; Leu = Leucocytes; Lym = Lymphocytes; Mon = Monocytes; N = Neutrophils; Eos = Eosinophils; Bas = Basophiles; Ery = Erythrocytes; MCV = Mean corpuscular volume; Hct = Hematocrit; MCH = Mean corpuscular haemoglobin content; MCHC = Mean corpuscular hemoglobin concentration; Hgb = Hemoglobin; Plt = Blood plate; N/Lym = Neutrophils/Lymphocyte ratio.
Superscripts refer to probability levels for significance tests ( <sup>\*</sup> *P* < 0.05, <sup>\*\*</sup> *P* < 0.01, <sup>\*\*\*</sup> *P* < 0.001)

Furthermore, the 213 animals with available haematological measurements at 28 d were analyzed to explore their association with the model parameters (Table 3). Parameter *ABC* was the parameter that showed the highest number of significant correlations, showing the strongest correlations with Hct (r= −0.38, *p*-value= 6.25 × 10^−9^), Hgb (r= −0.32, *p*-value= 2.57 × 10^−6^), MCHC (r= 0.32, *p*-value= 2.00 × 10^−6^), MCV (r= −0.32, *p*-value= 1.40 × 10^−6^), and Ery (r= ×0.20, *p*-value= 2.92 × 10^−3^). Regarding the parameter *D* it should be emphasized that it was significantly negatively associated with MCV (r= −0.22, *p*-value= 1.35 × 10^−3^) and Mon (r= −0.20, *p*-value= 3.46 × 10^−3^).

In the case of parameter *C*, the strongest negative significant associations were with Hct (r= −0.24, *p*-value= 4.05 × 10^−4^), MCV (r= −0.23, *p*-value= 5.71 × 10^−4^), and Hgb (r= −0.21, *p*-value= 1.90 × 10^−3^).

**Table 3.** Pearson’s coefficients to visualize correlations among the model parameters of the Gompertz-Makeham perturbed model and the haematological measurements (28 d) (N= 213 animals)

|  | *D* | *C* | *t<sub>s</sub>* | *ABC* | BW_w | Age_w | Leu | Lym | Mon | N | Eos | Bas | Ery | MCV | Hct | MCH | MCHC | Hgb | Plt | N/Lym |
| --- | --- | --- | --- | --- | --- | --- | --- | --- | --- | --- | --- | --- | --- | --- | --- | --- | --- | --- | --- | --- |
| $\mu_0$ | 0.88*** | 0.49*** | 0.41*** | 0.42*** | -0.18** | 0.30*** | 0.02 | 0.12 | -0.13 | -0.09 | 0.15* | -0.11 | 0.01 | -0.13 | -0.08 | -0.03 | 0.15* | -0.02 | -0.04 | -0.10 |
| *D* |  | 0.40*** | 0.33*** | 0.37*** | 0.07 | 0.36*** | 0.05 | 0.18** | -0.20** | -0.14* | 0.18** | -0.12 | -0.05 | -0.22** | -0.19** | -0.12 | 0.16* | -0.16* | 0.04 | -0.15* |
| *C* |  |  | 0.03 | 0.63*** | 0.11 | 0.24*** | -0.13 | 0.06 | -0.12 | -0.02 | 0.02 | -0.15* | -0.11 | -0.23*** | -0.24*** | -0.12 | 0.18** | -0.21** | 0.03 | -0.04 |
| *t<sub>s</sub>* |  |  |  | 0.36*** | -0.16* | 0.09 | -0.10 | 0.02 | -0.01 | -0.01 | 0.01 | -0.05 | 0.10 | -0.02 | 0.07 | 0.02 | 0.06 | 0.11 | 0.02 | -0.03 |
| *ABC* |  |  |  |  | 0.27*** | 0.39*** | -0.12 | 0.14* | -0.14* | -0.11 | 0.09 | -0.19** | -0.20** | -0.32*** | -0.38*** | -0.13 | 0.32*** | -0.32*** | 0.17* | -0.14* |

$\mu_0$ = individual growth rate at the moment of weaning; *D* = extent of the exponential decay of the growth; *C* = level of perturbation; *t<sub>s</sub>* = moment at which the animal recover for the perturbation; *ABC* = area between curves; BW\_w = body weight at weaning; Age\_w = age at weaning; Leu = Leucocytes; Lym = Lymphocytes; Mon = Monocytes; N = Neutrophils; Eos = Eosinophils; Bas = Basophiles; Ery = Erythrocytes; MCV = Mean corpuscular volume; Hct = Hematocrit; MCH = Mean corpuscular haemoglobin content; MCHC = Mean corpuscular hemoglobin concentration; Hgb = Hemoglobin; Plt = Blood plate; N/Lym = Neutrophils/Lymphocyte ratio.
Superscripts refer to probability levels for significance tests ( \* *P* < 0.05, \*\* *P* < 0.01, \*\*\* *P* < 0.001).

## Discussion

The challenge to obtain reliable estimates of weaning robustness in large populations motivated us to investigate the capability of a modelling approach from piglets’ growth curves. Here we describe how we achieve this objective by implementing a perturbed modelling approach of growth curves by using the Gompertz-Makeham function. This model successfully characterises growth resilience by using live weight, which could be recorded in swine production systems. This facilitates their future deployment in pig breeding systems. In addition, the implementation of the approach is made available via the SciLab open-source numerical computation package.

### The Gompertz-Makeham function efficiently models the growth perturbation at weaning

Previous studies evaluating different functions for describing growth in pigs (Schulin-Zeuthen *et al*., 2008) shown the bounties of the most used functions to represent changes in BW. Due to the practical implications of characterising growth in pigs, different models have been developed to represent growth dynamics (Schulin-Zeuthen *et al*., 2008).

To our knowledge, this is the first report in modelling the live weight response of piglets’ at weaning. Various statistical and reliability measures of the model are obtained related to the weaning perturbation, including parameters related to the depth, to the decay of the growth, and the time recovery derived from this perturbation. The application to the population data demonstrates that these parameters appear close to normal distribution. Our study suggests that this model development is instrumental to define parameters for evaluating the level of robustness of animals in current swine production systems.

### Applicability of the model in porcine production

Since more farms invest in precision livestock technologies, such as a weight collecting systems (Stygar *et al*., 2018), the opportunity of deploying modelling approaches into real-life situations becomes more feasible. Our study developed a mathematical model to describe piglets’ robustness at weaning by using the live weight during the first 75 days after weaning. We introduce here five weaning robustness parameters that have shown correlations to relevant haematological traits at weaning: *μ_0_, C, D, t_s_* and *ABC*. The first one, parameter *μ_0_*, characterise the individual growth rate at the moment of weaning. Parameter *C*, is a measure of the depth of perturbation. The practical use of parameter *C* as a measure of the perturbation relies on the idea that the deeper the perturbation, the higher the degree of sensitivity to the weaning stress of the animal. This parameter should be interpreted together with the parameter *D*, which indicates the extent of the exponential decay of the growth due to the perturbation. In this context, *D* reflects more growth robustness than resilience to the weaning perturbation. In the developed perturbed model, the animal response to the perturbation is partitioned into a perturbation and a recovery window, with the transition being defined by the parameter *t_s_*. As with *C*, the *t_s_* parameter would be related to the concept of resilience as a lower *t_s_* would represent an animal that is recovering faster from the perturbation.

In an attempt to create a measure that summarizes the previous reported model parameters we have proposed the *ABC* index, which reflects the difference between a theoretical unperturbed curve and the perturbed model response. We consider that this parameter is a good candidate to represent the global robustness of the animals at weaning, because it informs on the animal capabilities in terms of the amplitude and length of perturbation, and the rate of animal recovery. For a real implementation at large scale in pig breeding systems, using this *ABC* index has great advantages in terms that provides an indicator on the robustness/resilience to weaning. An alternative to explore could be the idea of developing a robust index integrating the different parameters of the perturbed model.

It should be note that modelling approach here developed is built on the basis that the BW animal measurements are frequently recorded, with particular attention to the first weeks after weaning. The limitation of our approach is that to guarantee a robust model, high frequency of animal measurements is required. Nevertheless, this limiting factor should be minimised with the generalisation of modern farm systems with automatic measurement recording.

### The proposed robustness parameters correlate with diarrhoea score and blood formula

A positive correlation between parameter *C* and the faeces score data was found in the present study. Given that the faeces score is a measure of the prevalence of digestive disorders, this indicates that the perturbed model provides useful information on the health component of robustness to weaning. Weaning is usually associated with a dramatic reduction in food intake, resulting in altered structure of the small intestine. Piglets usually showed weight loss during the first 3 days post weaning. During the first week post weaning, the reported prevalence of diarrhoea in at least one day is 73% (Vente-Spreeuwenberg *et al*., 2003).

Blood constitutes a relevant tissue for phenotyping immune capacity (Flori *et al*., 2011; Mach *et al*., 2013; Schroyen and Tuggle, 2015), and evaluating the health status of pigs. To study the health status in piglets during the period after weaning, we conducted correlation analyses between the haematological traits collected at weaning (28 d) and one week after weaning (34 d), and the model parameter data. The first week after weaning is considered the most stressful period for piglets, when intestinal dysfunction and changes in the metabolism occur (Campbell *et al*., 2013), as well as changes in physiological and immunological parameters (Kick *et al*., 2012; Pié *et al*., 2014).

With respect to the blood measurements made at weaning (28 d), a moderate positive correlation between the parameter *ABC* and the level of MCHC (r= 0.32, *p*-value= 2.00 × 10^−6^), and a negative correlation with MCV levels (r= −0.32, *p*-value= 1.40 × 10^−6^) was found. Some studies have suggested that MCHC and MCV parameters are early indicators of iron deficiency (Svodoba *et al*., 2008). Additionally, negative correlations were found between this *ABC* parameter and the percentage of Hct (r= −0.38, *p*-value= 6.25 × 10^−9^) and the levels of Hgb (r= −0.32, *p*-value= 2.57 × 10^−6^) at 28 days of age. It has been demonstrated that there is a positive association between Hgb, Hct and average daily gain in the three weeks post-weaning period (Bhattarai and Nielsen, 2015). These authors also reported that an increase in 10 g haemoglobin/l blood corresponded to a weight gain improvement of 17.2 g daily weight gain. The negative correlations observed in this study between the parameter *ABC* and Hgb and Hct are in concordance with the reported publication, showing that those perturbed animals grow less.

Regarding the blood measurements collected one week after the weaning period (34 d), the negative correlation identified between parameter *ABC* and the percentage of Mon (r= −0.30, *p*-value= 5.15 × 10^−8^) is noteworthy. The levels of Mon could be a good estimator of the animal health status due to its important roles in both innate and adaptive immune responses, killing microbial pathogens and tumour cells, and exerting immunoregulatory functions through cytokine production and processing and presentation of antigens to Lymphocytes (Chamorro *et al*., 2005). Moreover, it is assumed that pigs are physiologically and immunologically competent by 35 d of age (Kick *et al*., 2012).

Whilst promising, the interpretation of the correlations between the proposed resilience and robustness indicators (derived from the growth curve models) and the health status measurements should be interpreted with caution until they can be tested across a broader range of herds.

## Conclusions

This study presents a method to quantify and define parameters related to piglet weaning robustness. These parameters are derived by modelling piglet body weight trajectories from weaning onwards. This work provides biological parameters that inform on the amplitude and length of perturbation, and the rate of animal recovery. In addition, we have identified significant correlations between the model parameters, and individual diarrhoea scores and haematological measurements, which illustrate the usefulness of these parameters as potential components of an integrated robustness index.

## Acknowledgements

We are grateful to the personal at GENESI’s farm at *Le Magneraud* for their implication on the generation of animals and samples. Experiments were funded by the Pigletbiota project by the French *Agence Nationale de Recherche* (ANR; project: ANR-14-CE18-0004). We are grateful to all members of the Pigletbiota consortium that support this project, and which include DELTAVIT (CCPA group), InVivo-NSA (InVivo group), LALLEMAND, SANDERS (AVRIL group), and TECHNA companies and the ALLIANCE R&D association (AXIOM, CHOICE GENETICS, NUCLEUS and IFIP). We also grateful to the Valorial competitiveness cluster for its support to the project.

## Ethics statement

All animal procedures were performed according to the guidelines for the care and use of experimental animals established by INRA (Ability for animal experimentation to J. Estellé: R-94ENVA-F1-12; agreement for experimentation of INRA’s *Le Magneraud* farm: A17661; protocol approved by the French Ministry of Science with authorization ID APAFIS#2073-2015092310063992 v2 after the review of ethics committee n°084).

## Software and data repository resources

The Scilab code for calibration of the perturbed model is available under request for academic purposes.

## Supplementary material

**Supplementary Table S1:** Parameter estimates of the perturbed model and comparison between Gompertz and perturbed models performances

